# Mapping functional regions of essential bacterial proteins with dominant-negative protein fragments

**DOI:** 10.1101/2022.01.04.474984

**Authors:** Andrew Savinov, Andres Fernandez, Stanley Fields

## Abstract

Massively-parallel measurements of dominant negative inhibition by protein fragments have been used to map protein interaction sites and discover peptide inhibitors. However, the underlying principles governing fragment-based inhibition have thus far remained unclear. Here, we adapt a high-throughput inhibitory fragment assay for use in *Escherichia coli*, applying it to a set of ten essential proteins. This approach yielded single amino acid-resolution maps of inhibitory activity, with peaks localized to functionally important interaction sites, including oligomerization interfaces and folding contacts. Leveraging these data, we perform a systematic analysis to uncover principles of fragment-based inhibition. We determine a robust negative correlation between susceptibility to inhibition and cellular protein concentration, demonstrating that inhibitory fragments likely act primarily by titrating native protein interactions. We also characterize a series of trade-offs related to fragment length, showing that shorter peptides allow higher-resolution mapping but suffer from lower inhibitory activity. We employ an unsupervised statistical analysis to show that the inhibitory activities of protein fragments are largely driven not by generic properties such as charge, hydrophobicity, and secondary structure, but by the more specific characteristics of their bespoke macromolecular interactions. AlphaFold computational modeling of peptide complexes with one protein shows that the inhibitory activity of peptides is associated with their predicted ability to form native-like interactions. Overall, this work demonstrates fundamental characteristics of inhibitory protein fragment function and provides a foundation for understanding and controlling protein interactions *in vivo*.

**Significance Statement:** Peptide fragments derived from protein sequences can inhibit interactions of their parental proteins, providing a promising avenue for drug development. Here we employ a massively-parallel assay to measure *in vivo* inhibition by fragments that tile the full sequences of ten essential bacterial proteins. We leverage these data to decipher principles of fragment-based inhibition, showing how parental protein concentration drives activity and how protein fragment length interplays with activity and specificity. We employ statistical analysis to parse the roles of biophysical properties in fragment-to-fragment variation, and AlphaFold modeling to determine the relationship between measured inhibitory activity and predicted native-like binding. These results provide a path towards rational design of peptide inhibitors and broader principles of protein-protein interactions in living cells.

## Introduction

Peptides have the capability to act as potent modulators of biological function by binding to proteins at specific sites, thereby altering protein activity. Compared to small molecules, peptides have advantages in specificity, ability to target large and shallow interaction surfaces (1, 2), and genetic encodability. Organisms have leveraged this phenomenon by encoding peptide antimicrobials (3) as well as biological peptides termed small proteins or miniproteins, which frequently regulate larger proteins (4–7). Synthetic peptides can also be engineered for such inhibitory functions, providing potential avenues for novel cancer therapeutics (8), antivirals (9) and antibiotics (10).

Fragments of native protein sequences, in particular, can reproduce native interactions with binding partners while lacking other functions of the full-length protein. Similar to the case with truncation mutants (11), these binding events allow such peptides to compete with their parental protein for its native interactions, and thereby act as dominant negative inhibitors. While most such examples of dominant negative inhibition involve intermolecular interactions, protein fragments can also inhibit intramolecular interactions involved in protein folding (12, 13). Given their interference with native binding contacts, inhibitory fragments permit functional mapping of proteins, identifying important interaction sites at sub-gene resolution. DNA sequencing-based methods allow the identification of such peptides in a massively-parallel manner (14–16) to map functional domains (14, 15). In these assays, a growth selection is performed on cells carrying a protein fragment library under conditions in which the parental protein is important for growth. Depletion of fragment-encoding sequences from the population, reflecting inhibitory activity, is quantified by performing high-throughput sequencing before and after selection. Peptides derived from such approaches are capable of pulling down their expected interaction partners (15).

Here, we adapted this method to map a diverse set of essential bacterial proteins in *E. coli*, reasoning that the abundance of structural and functional information available for this organism should reveal biophysical principles underlying fragment-based inhibition as well as identify possible target sites for developing antimicrobials. These measurements permitted functional mapping at single residue resolution, revealing the importance of dozens of structural elements that can act as dominant-negative peptides. Many inhibitory protein fragments mapped to protein-protein interaction sites, and others mapped to regions that inhibit intramolecular folding interactions. Leveraging the data across diverse proteins, we performed analyses that revealed the importance of fragment length, as well as a strong negative correlation between the cellular concentrations of proteins and their susceptibility to fragment-based inhibition. We also employed ANOVA analysis and AlphaFold modeling to delve into the roles of biophysical properties in the inhibitory activity of fragments. In sum, our results provide key principles for understanding protein fragment inhibition in living cells.

## Results

### A high-throughput protein fragment assay recapitulates a folding-inhibitory region of dihydrofolate reductase

To scan for dominant-negative inhibitory fragments of *E. coli* proteins, we developed a high-throughput assay (Fig 1*A*; *Materials and Methods*) analogous to approaches employed in yeast and human cells (14, 15, 17). We combined measurements of growth selection depletion with an alignment of fragments to their parental proteins, generating maps of inhibitory activity as a function of sequence position. To systematically cover all positions of a protein with fragments of tunable length, we array-synthesized libraries of DNA fragments tiling across the coding sequence of each protein with single residue resolution (*Materials and Methods*), similar to the approach of Ford *et al.* (15).

**Figure 1:**
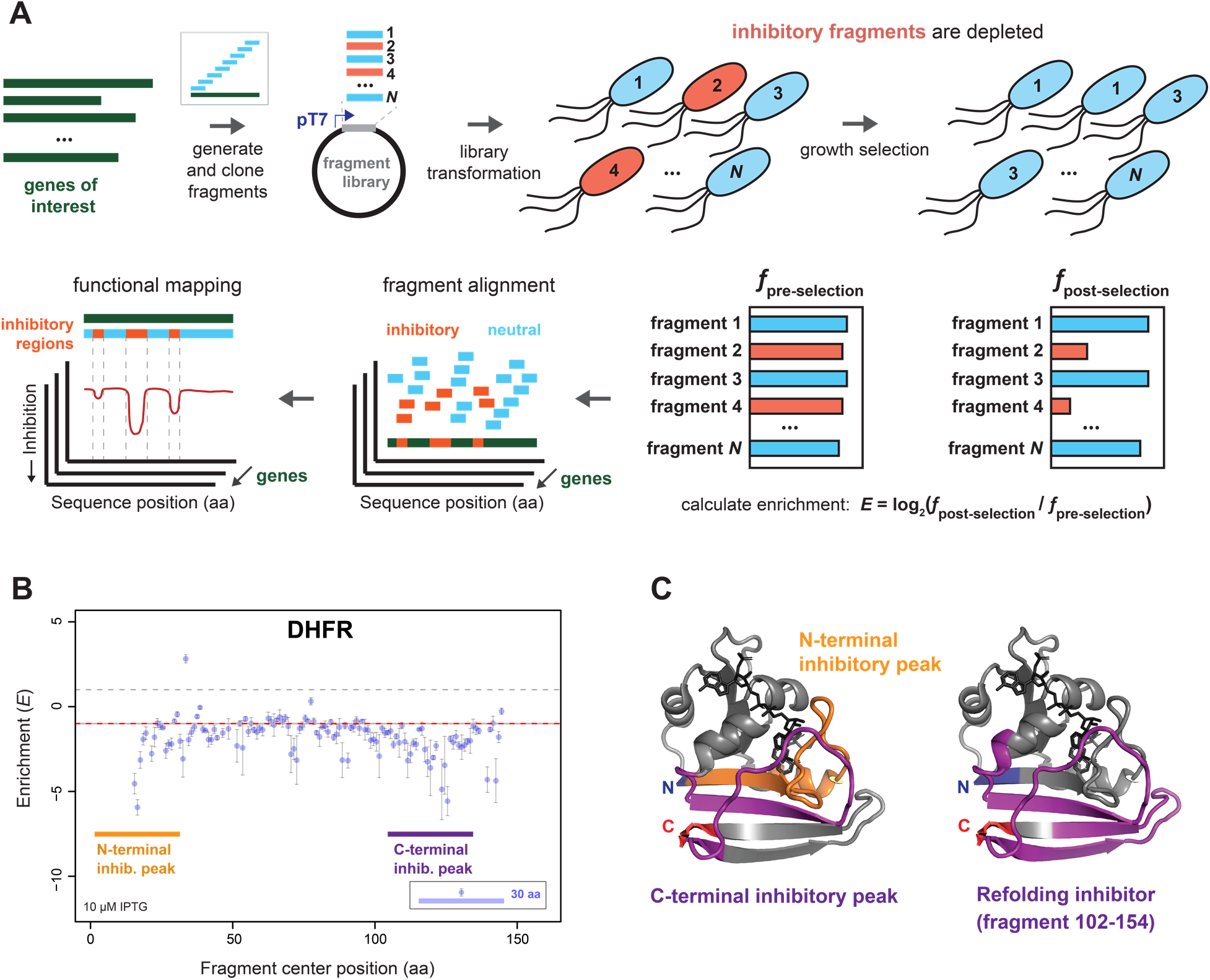
Mapping protein functional regions with dominant negative inhibitory fragments. (*A*) Schematic describing the high-throughput inhibitory fragment assay. Fragments generated from genes of interest are cloned into an expression vector and transformed into *E. coli*. A growth selection is performed and inhibitory fragment depletion from the population is quantified by high-throughput sequencing, allowing determination of the enrichment *E* = log2(*f*post-selection/*f*pre-selection), where *f* denotes frequency, with *E* < 0 indicating depletion. Fragment alignment to the parental protein sequence allows identification of inhibitory regions. (*B*) Tiling inhibitory fragment scan results for *E. coli* DHFR, at a tile step size of 1 amino acid. Enrichment (*E*) is plotted as a function of fragment center position. Points: individual fragment measurements; error bars: s.e.m. from multiple experiments; dashed horizontal lines: guide lines at ±1*E* (±2-fold change in frequency). Colored rectangular markers indicate positions of inhibitory peaks, with width indicating a representative fragment. (*C*) DHFR crystallographic structure (PDB ID 7dfr) with N- and C-termini indicated, overlaid by the N-terminal and C-terminal inhibitory peaks (*left*), and the known refolding-inhibitory fragment (*right*) from the work of Hall and Frieden (12).

We sought to establish this approach in *E. coli* using a model protein with a known inhibitory fragment acting by a well-characterized mechanism: dihydrofolate reductase (DHFR), a monomeric protein for which a C-terminal fragment inhibits the refolding of the full-length protein *in vitro* (12). We reasoned that similar peptide fragments might also affect DHFR folding *in vivo*. We generated an inhibitory fragment map by plotting the inhibitory activity (selection enrichment, *E*) of 30-residue fragments as a function of sequence position (Fig. 1*B*). We observed an inhibitory peak in the same C-terminal region of the protein as the known folding inhibitor (12) (Fig. 1*C*), suggesting that these peptides also inhibit folding in cells. We additionally observed an N-terminal inhibitory peak, mapping to a region involved in binding dihydrofolate, that might inhibit DHFR via substrate titration (Fig. 1*C*). We used a Nextera transposase-based random fragment library (14) of the DHFR-encoding *folA* gene to demonstrate coding frame-dependence of inhibition, with out-of-frame fragments predominantly neutral (*E* ≈ 0) (Fig. S1); these results underscored that fragment inhibitory activity is sequence-dependent and acts at the protein level.

### Tiling fragment scans of essential *E. coli* proteins map key interaction sites

Given the results with DHFR, we applied high-throughput inhibitory fragment mapping to a set of nine additional bacterial proteins (Table 1), spanning several core processes: DNA replication (GyrA, Ssb), transcription (RpoB), translation (IleS, RplL), protein quality control (GroEL, GroES), cell division (FtsZ), and outer membrane maintenance (LptG). Overall, this assay yielded a raw enrichment signal that delineated inhibitory peaks without requiring coverage-based smoothing (Fig. 2*A-G*) as employed previously (14, 15). An overview of the results is presented in Tables 1 and 2.

**Figure 2:**
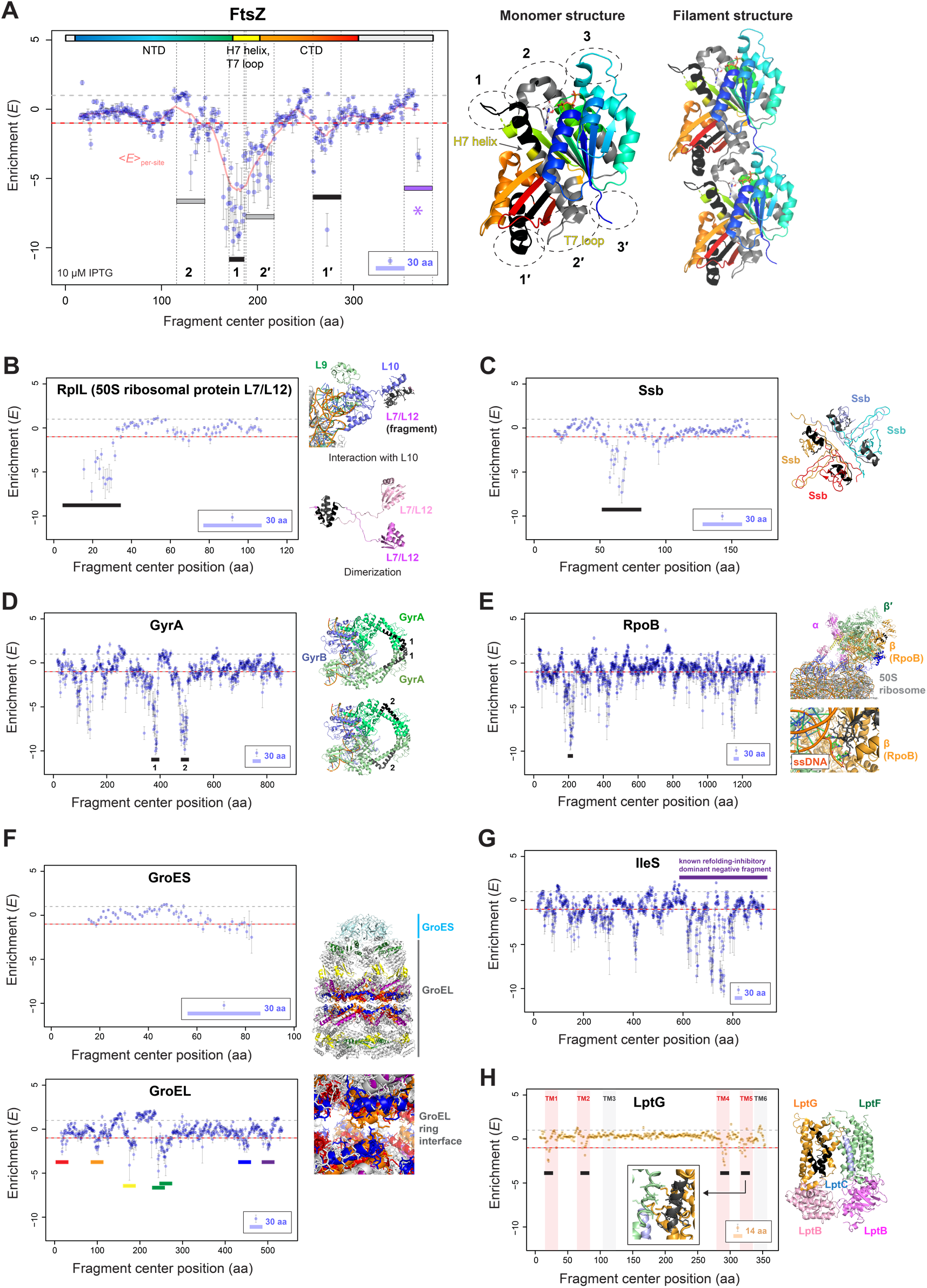
Inhibitory fragments map interaction interfaces of essential bacterial proteins. Inhibitory fragment scans of *E. coli* proteins with tiling fragments, at a tile step size of 1 amino acid. Enrichment (*E*) is plotted as a function of fragment center position. Points: individual fragment measurements; error bars: s.e.m. from multiple experiments; dashed horizontal lines: guide lines at ±1*E* (±2-fold change in frequency). Colored rectangular markers indicate positions of inhibitory peaks (with width indicating a representative fragment, unless otherwise noted). In protein structures to the right of the plots, locations of inhibitory peaks (generally representative fragments) are overlaid on the structure in a color matching the corresponding peak marker in the plot to the left (generally black). (*A*) – (*G*) 30-residue tiling fragment scans of indicated proteins; (*H*) 14-residue tiling fragment scan of LptG. Additional features are present as indicated: (*A*), salmon line indicates average *E* per residue due to all fragments covering that residue; the width of the peak **1** marker indicates the peak width (16 residues); the lavender asterisk marks the inhibitory peak that falls in the C-terminal intrinsically disordered region; and the colored bar at the top of the plot indicates the layout of structural elements in the linear sequence; (*H*), shaded regions denote indicated topological domains. PDB IDs for structures: (*A*) FtsZ: 6unx; (*B*) RplL: 3j7z (*upper*) and 1rqu (*lower*); (*C*) Ssb: 1eqq; (*D*) GyrA: 6rks; (*E*) RpoB: 6x9q; (*F*) GroEL-ES: 1aon; (*H*) LptG: 6mi7. No structure is shown for *E. coli* IleS because it has not yet been determined.

**Table 1:** Overview of inhibitory fragment assay targets, their properties, and overall results. Properties of each *E. coli* protein assayed by inhibitory fragment scanning are compiled, together with the average inhibitory effects of tiling 30-residue fragments. Cellular concentrations of proteins are from ref. 24.

| *E. coli* gene | Protein | Length (aa) | Cellular abundance (protein molecules/cell) | <E> for 30aa tiling fragments |
| --- | --- | --- | --- | --- |
| *folA* | Dihydrofolate reductase (DHFR) | 159 | 2,515 | -1.78 |
| *gyrA* | DNA gyrase subunit A (GyrA) | 875 | 5,219 | -1.21 |
| *ileS* | Isoleucyl tRNA synthetase (IleS) | 938 | 6,143 | -1.26 |
| *ftsZ* | Cell division protein FtsZ | 383 | 6,750 | -1.20 |
| *rpoB* | RNA polymerase, $\beta$ subunit (RpoB) | 239 | 16,156 | -0.99 |
| *rplL* | 50S ribosome protein L7/L12 (RplL) | 1342 | 472,965 | -0.84 |
| *ssb* | Single-stranded DNA binding protein (Ssb) | 121 | 14,444 | -0.66 |
| *groL* | 60 kDa chaperonin (GroEL) | 178 | 52,165 | -0.28 |
| *groS* | 10 kDa chaperonin (GroES) | 548 | 59,001 | -0.21 |
| *lptG* | Lipopolysaccharide export system permease protein LptG | 97 | 775 | — |

We first consider measurements with tiling fragments of 30 residues, as used with DHFR. The results for FtsZ (Fig. 2*A*, *left*) exemplify what can be learned using this approach. FtsZ forms filaments that are essential for bacterial cell division (18); monomers polymerize head-to-tail (19), providing a well-characterized binding interface (Fig. 2A, *right*). Several binding contacts from either side of this interface yielded inhibitory fragment peaks. The two strongest inhibitory peaks map to loop structures with flanking helices, which form portions of the oligomerization interface that we term sites **1** and **2′** (Fig. 2*A*); inhibitory fragment peaks also mapped to the complementary regions of the adjacent FtsZ monomer, sites **1′** and **2** (Fig. 2*A*). Conversely, no inhibitory peaks were observed mapping to interaction interface regions **3** and **3′** (residues 65-69 and 1-11; Fig. 2*A*), consistent with a B-factor analysis suggesting that the interactions of these sites are weak (19). Outside of the head-to-tail monomer interface, the fragment scan detected an inhibitory region at the junction of the intrinsically disordered C-terminal linker and the C-terminal-most 15 residues, which enable modulation of FtsZ activity by binding partners such as FtsA, ZipA, and MinC (20). This result demonstrates the utility of inhibitory fragment scanning for functionally mapping intrinsically disordered regions for which no structural information is available.

Inhibitory fragments mapped to key interaction sites across the sequences of several additional proteins. For the 50S ribosomal subunit protein L7/L12 (RplL), we observed a single inhibitory peak localized to an N-terminal region responsible for RplL dimerization and binding to 50S ribosomal protein L10; both interactions are required to form the ribosomal stalk (Fig. 2*B*). For the single-stranded DNA binding protein (Ssb), the single inhibitory peak maps to an alpha-helical region involved in the dimerization of Ssb, which further tetramerizes (Fig. 2*C*). For DNA gyrase A (GyrA), multiple inhibitory peaks were observed (Fig. 2*D*), with the strongest two mapping to the C-Gate formed by a GyrA dimer interface; the C-Gate must properly open and close to control the directionality of DNA strand transfer during gyrase activity, with errors resulting in double-stranded DNA breaks (21). Several peaks were also identified for the beta subunit of RNA polymerase (RpoB), including one localized to a site that interacts with single-stranded DNA in the context of a transcription-translation “expressome” complex (Fig. 2*E*; ref. (22)). For GroEL, several inhibitory fragment peaks map to the interface between the two stacked GroEL heptamers (Fig. 2*F*, *right* and *bottom left*), suggesting inhibition of proper GroEL/ES complex assembly and chaperone function. On the other hand, we observed no inhibitory peaks in a fragment scan of GroES (Fig. 2*F*, *top left*).

For isoleucyl tRNA synthetase (IleS), the strongest inhibitory peaks cluster in the C-terminal region (Fig. 2*G*). The structure of *E. coli* IleS has not been determined, but these results are congruent with functional data; the strong C-terminal inhibitory peaks overlap with a large C-terminal fragment (residues 585-939) that acts as a folding inhibitor *in vitro* and a dominant negative *in vivo* (13). The results of the inhibitory fragment scan suggest that this folding-inhibitory activity is localized more finely in the sequence, with several distinct peaks at residues ∼600-800.

To determine whether membrane proteins are amenable to fragment-based inhibition, we performed an inhibitory fragment scan of LptG (lipopolysaccharide transport protein G), a transmembrane protein involved in lipopolysaccharide transport from the inner to the outer membrane. To better resolve functions of the multiple short cytosolic, transmembrane, and periplasmic domains found throughout the sequence, we scanned LptG with fragments of 14 residues. The fragment scan yielded four inhibitory peaks, which all map to transmembrane alpha-helices (TM1, TM2, TM4, or TM5; Fig. 2*H*). It seems unlikely that this inhibition arises from mechanisms generic to hydrophobic peptides, *e.g.*, aggregation, as fragments mapping to the remaining two transmembrane alpha-helices, TM3 and TM6, did not yield inhibitory peaks. Rather, the inhibitory fragments likely localize to the membrane and inhibit LptG folding and/or interactions with other membrane proteins. For example, cryo-EM structures of the Lpt complex (23) reveal that LptG TM5 forms a binding interaction with the TM1 helix of LptF. Similarly, LptG TM1 interacts with the alpha-helical protein LptC (23), although the inhibitory fragment peak maps to the center of the TM1 helix rather than the end that directly binds LptC.

### A simple principle underlies variable susceptibility of proteins to fragment-based inhibition

We sought to understand the factors driving the varied susceptibility of different proteins to fragment-based inhibition. For example, fragments derived from GroEL and especially GroES had lower average inhibitory activity than fragments derived from the other *E. coli* proteins (Fig. 2*F, left* and Table 1). In contrast, fragments derived from DHFR exhibited a baseline inhibitory effect (Fig. 1*B*), resulting in DHFR fragments exhibiting the highest average inhibitory activity of the proteins tested (Table 1). Many factors could play a role in this differential susceptibility, including localization, folding stability, structural characteristics, number of interaction partners, strength of native binding interactions, and cellular concentration of the parental proteins. This last factor seemed likely to be important, as a given concentration of fragments that bind competitively with native interactions (including folding interactions) should be less inhibitory when the parental protein is more abundant.

We performed an analysis of the relationship between the average inhibitory effects of 30-residue fragments derived from each protein and the corresponding protein concentration (protein copy number per cell; refs. (24, 25)). We uncovered a robust negative correlation between parental protein concentration and susceptibility to fragment-based inhibition, observed as a linear relationship (R^2^ = 0.95) between the log of the protein copy number per cell and the average enrichment <*E*> of fragments derived from that protein (Fig. 3), with a higher <*E*> corresponding to lower average inhibitory activity. The slope indicates that a ∼10-fold increase in protein concentration is associated with ∼50% less fragment depletion in our assay (an average *E* increase of ∼1). Thus, the poor susceptibility of GroEL and GroES to fragment-based inhibition might be explained by their high cellular concentrations, and the broad sensitivity of DHFR to fragment-based inhibition by its low concentration (Fig. 3, Table 1). The sole exception to the correlation was the ribosomal protein RpIL, fragments of which were much more inhibitory than expected from its cellular concentration (Fig. 3). A possible reason for this exception is that even weak inhibition of ribosome assembly may be amplified in the assay due to the large effect of ribosome levels on growth rates (26).

**Figure 3:**
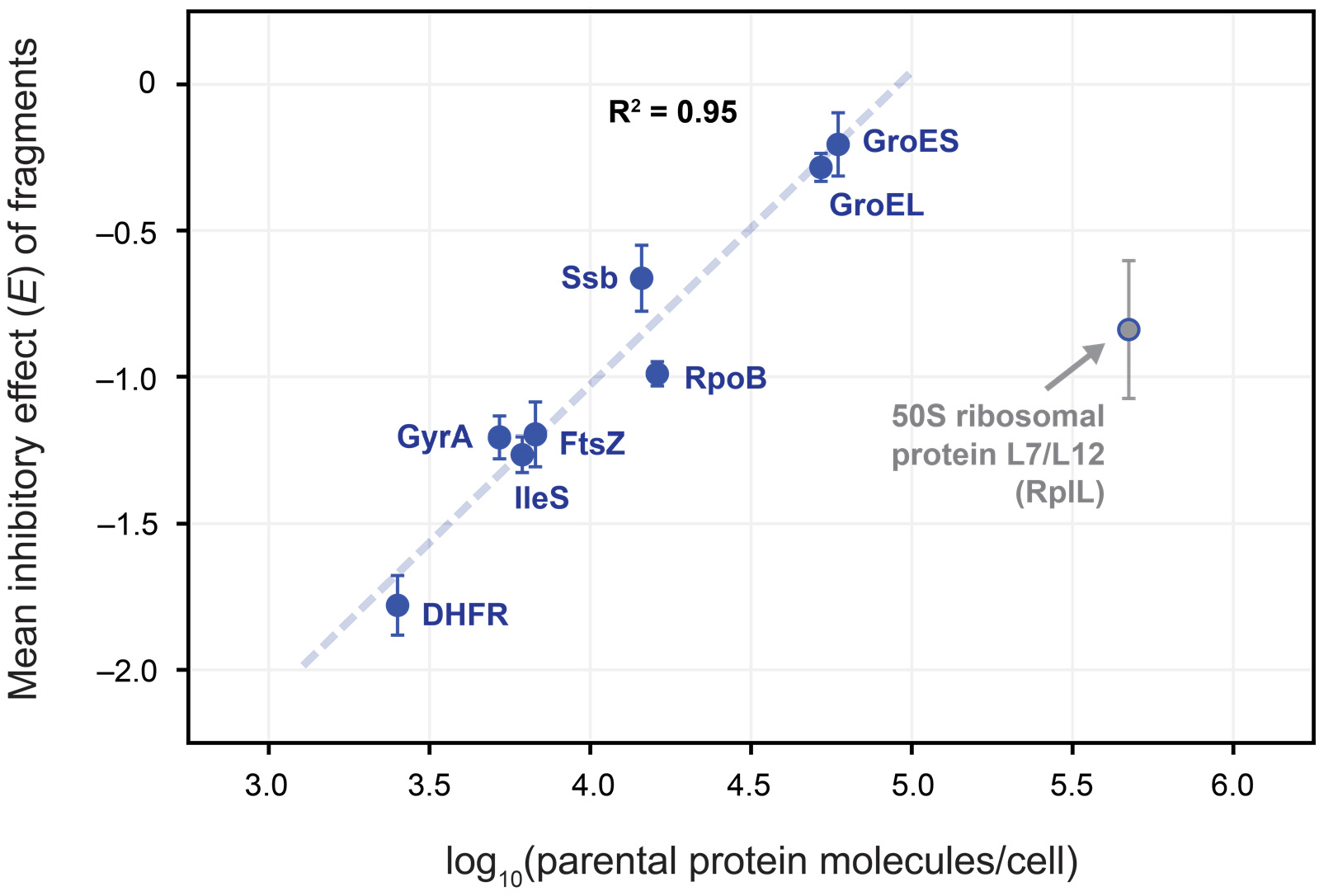
Cellular protein concentration negatively correlates with average susceptibility to fragment-based inhibition. Plot of mean inhibitory effects of 30-residue fragments tiling each of the cytoplasmic proteins included in this study as a function of the log of the cellular concentration of the full-length protein, from Li *et al.* (24). The 50S ribosomal protein L7/L12 outlier is indicated. Error bars: s.e.m. A linear fit is shown to the data for the non-ribosomal proteins.

In sum, these results suggest that protein fragment-based inhibition is driven primarily by fragments acting as competitive inhibitors of native interactions formed by their parental proteins, despite the complexities of the *in vivo* scenario. Additionally, these findings are inconsistent with the idea that the inhibitory effects of fragments are largely due to more generic effects, such as toxicity due to peptide aggregation.

### Relationship between fragment length and inhibitory activity

We sought to understand how the inhibitory effects of protein fragments might depend on fragment length. For example, shorter fragments might allow finer mapping of structural elements, but peptides that are too short might have little effect due to limited binding energy. Conversely, longer fragments might exhibit stronger inhibitory activity by forming larger interaction surfaces or folding more stably in isolation. To investigate these questions, we examined the results of scans performed with fragments of 20 or 50 residues, compared with those performed with 30-residue fragments (Fig. 4*A-C*, Fig. S2).

**Figure 4:**
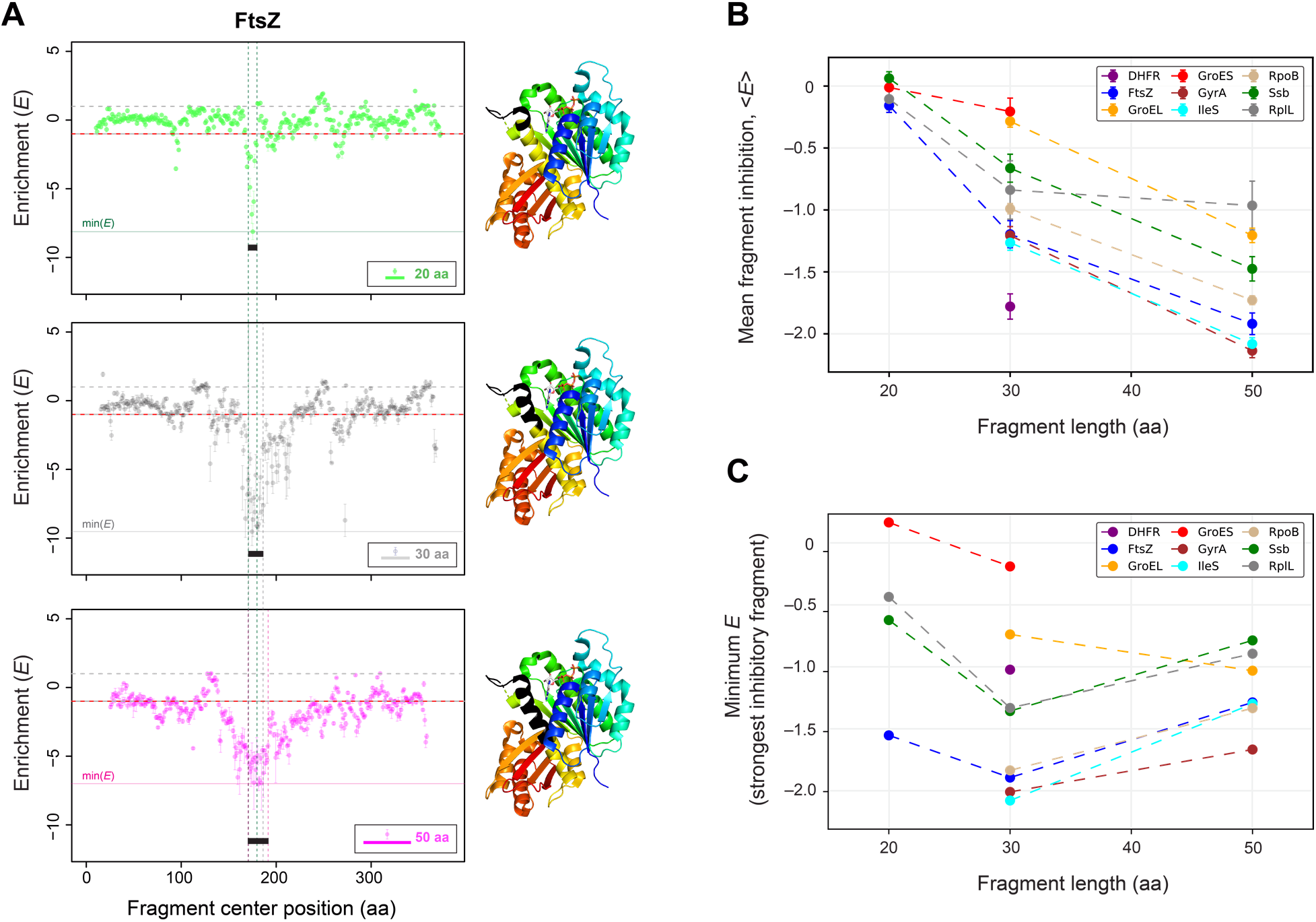
Inhibitory activity depends on fragment length. (A) Inhibitory fragment scans of FtsZ with tiling fragments, at a tile step size of 1 amino acid, comparing different fragment lengths. Data are plotted as in Figure 2, with individual fragment measurements and their s.e.m. across multiple experiments shown. A comparison of 20- (*top*), 30- (*middle*), and 50-residue (*bottom*) fragment scans is displayed. For each plot, the width of inhibitory peak **1** is indicated. Structures of FtsZ monomers to the right are overlaid with the width of the corresponding inhibitory peak in black. (*B*) Average inhibitory activity (*E*) of fragments tiling each cytosolic protein plotted as a function of fragment length. (*C*) Strongest inhibitory fragment among fragments tiling each protein, as a function of fragment length.

The results demonstrate several general features of fragment-based inhibition. First, tiling fragments of different lengths generally mapped out the same structural features of the parental proteins (Fig. 4*A*, Fig. S2). That the functional maps produced with different fragment lengths were so correlated – along with the finding that average inhibitory effects of fragments of 50 residues also negatively correlated with the concentrations of their parental proteins (Fig. S3) – further supports the idea that protein fragments typically act in a target- and sequence-specific manner.

Second, the average inhibitory activity was larger for longer fragments (Fig. 4*B*, Fig. S2). Increasing the fragment length from 20 to 30 residues increased the magnitude of inhibitory peaks while maintaining similar levels of background inhibition throughout the sequence (*i.e.*, the baseline was almost unaffected), representing an increase in both inhibitory activity and sequence-specificity. However, further increasing the length to 50 residues produced diminishing returns, as these fragments exhibited a marked increase in generic inhibitory activity across the sequence (a baseline shift to lower *E*), and, moreover, a decrease in the strength of many inhibitory peaks relative to the 30-residue fragments (*e.g.*, in FtsZ, RplL, Ssb, GyrA, IleS; Fig. 4*A*, Fig. S2A-D,H), which in almost all cases led to a decreased maximum inhibitory activity (higher minimum *E*) (Fig. 4*C*). We interpret this result as reflecting a competition between distinct target-specific and nonspecific inhibitory mechanisms (unrelated to perturbation of parental protein interactions) of the longer fragments, leading to reduced inhibition by 50-residue fragments from strong interaction sites. These results suggest that there is a fragment length between 20 and 50 residues that optimizes the balance of inhibitory activity and specificity; a length of 30 residues appears to be near this optimum (Fig. 4*C*).

Third, shorter fragments allowed for finer-resolution functional mapping, localizing inhibitory peaks to a narrower range of fragment center positions. In the case of FtsZ (Fig. 4*A*), the 20-residue fragment scan yielded a peak width of just 10 residues for the major inhibitory region (compared to 16 with 30-residue fragments, and 22 with 50-residue fragments), containing only one turn of the H7 alpha helix but still encompassing the loop of monomer interaction site **1** (Fig. 4*A*, *middle*). However, the weaker inhibitory activity on average also meant that some peaks observed with 30-residue fragments were less evident in the 20-residue fragment scan.

In sum, there are trade-offs between fragment length and important functional parameters that include inhibitory strength, specificity, and resolution of functional mapping. Shorter protein fragments allow higher-resolution mapping, but are associated with weaker inhibitory activity. Longer fragments can provide greater sequence-specific activity, but only to a point, with nonspecific background inhibition and reduced inhibitory peak magnitudes becoming evident by a fragment size of 50 residues in *E. coli*.

### The bulk of the variability in the inhibitory effects of individual fragments is case-specific

The effects of parental protein concentration and fragment length explain only a portion of the individual fragment-to-fragment variability (Fig. S4). Indeed, the strength of inhibition attainable at any specific site in a protein sequence must depend on unique features of each protein’s binding interactions. Hence, we sought to identify common features of individual fragments that might explain which sites in protein sequences are susceptible to fragment-based inhibition. Using analysis of variance (ANOVA), we approached this question across all protein targets investigated and several physicochemical and structural properties. In particular, we determined the percentage of total variability in the inhibitory effects of protein fragments that is attributable to hydrophobicity, charge, secondary structure features, fragment length, parental protein, relative position in the protein sequence, predicted peptide stability, and fragment-specific effects not explained by these other properties (see *Materials and Methods*; Table S1).

We found that a striking ∼72% of the total variability was attributable to fragment-to-fragment variation, outside of that attributable to any of the properties considered (Fig. 5). The properties that contributed the most to variability were fragment length, accounting for ∼10% of the total variation, and parental protein, accounting for ∼4.2%, matching the overall trends in fragment function. All other properties considered each explained less than 1% of the variation. Added together, secondary structure features of the fragment-covered portion of the protein structure explained ∼1.5% of the variation: ∼0.3-0.7% each for alpha-helical, beta-sheet, or turn content. Hydrophobicity had a comparable effect to secondary structure (∼0.7% of the variation) and charge an even smaller effect, well below that attributable to error, providing additional evidence against a generic inhibitory mechanism of aggregation due to charge and/or hydrophobicity, similar to the results of Ford *et al.* (15). Relative position of the fragment in the protein sequence and predicted peptide stability (27) were both negligible for explaining variability, each being associated with a fraction of the variation similar to or below the contribution of error.

**Figure 5:**
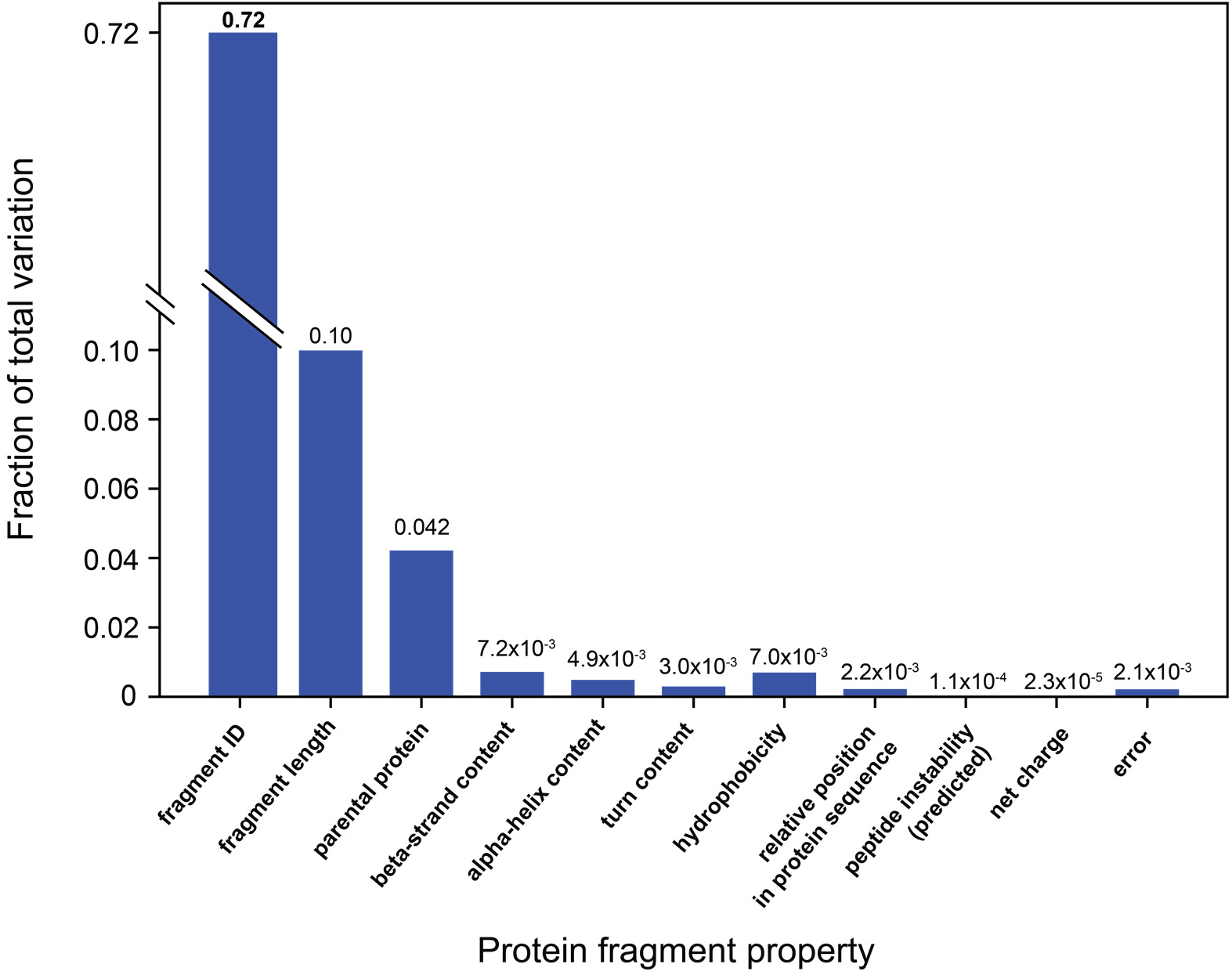
Generic properties are not the main drivers of protein-fragment-based inhibition. Fraction of total variation attributable to each property is plotted, based on a nested ANOVA. Properties include fragment length, parental protein, and secondary structure features of the fragment-covered sequence region of the full-length protein; “fragment ID” designates fragment-specific effects not attributable to variation in any of the properties considered.

The results of this analysis speak to the diversity of inhibitory function seen among protein fragments, mirroring the vast diversity of possible folds and interactions of proteins that provide the likely basis for most inhibitory activity.

### AlphaFold predictions of protein-peptide interactions provide a complementary approach

Our ability to predict protein fragments that inhibit native interactions might be improved by employing computational models of protein-peptide interactions. In particular, predicted modes of peptide binding and the extent to which they are native-like may correlate with the experimentally measured inhibitory strength of peptides from different binding interfaces. If so, peptide and interface properties used by the model to make these predictions might be extracted to determine design principles for inhibitory peptides. Given that, in the case of protein fragments, the major inhibitory mechanism appears to be the mimicry of native interactions, we reasoned that machine learning-based structure prediction algorithms trained on native folds and sequences would be a good choice of model.

We performed AlphaFold modeling of protein-peptide interactions (ref. (28); *Materials and Methods*), using the FtsZ results as a case study. This modeling predicted that representative peptides from the strongest inhibitory peak (site **1**) and its reciprocal peak (site **1′**) both form native-like interactions with FtsZ (Fig. 6, *top right*). Peptides derived from the second-strongest inhibitory peak (site **2′**) and reciprocal site **2** were predicted to form interactions with FtsZ that were non-native but nearby to their native interaction sites in the crystal structure, and still along the monomer interaction interface (Fig. 6, *middle right*). Finally, a peptide representing site **3′**, which yielded no inhibitory peak, was predicted to make only tenuous (and non-native) interactions with FtsZ (Fig. 6, *bottom right*), and a peptide from the similarly non-inhibitory site **3** was predicted to form substantial but non-native interactions, again along the interaction interface (Fig. 6, *bottom right*). These results thus show an approximate association between the potency of inhibitory protein fragment peaks and AlphaFold predictions of native-like binding for this case study, suggesting that these predictions indeed contain latent information on features underlying peptide-based inhibition that might be extracted to determine unifying properties of inhibitory fragments and enable fragment design.

**Figure 6:**
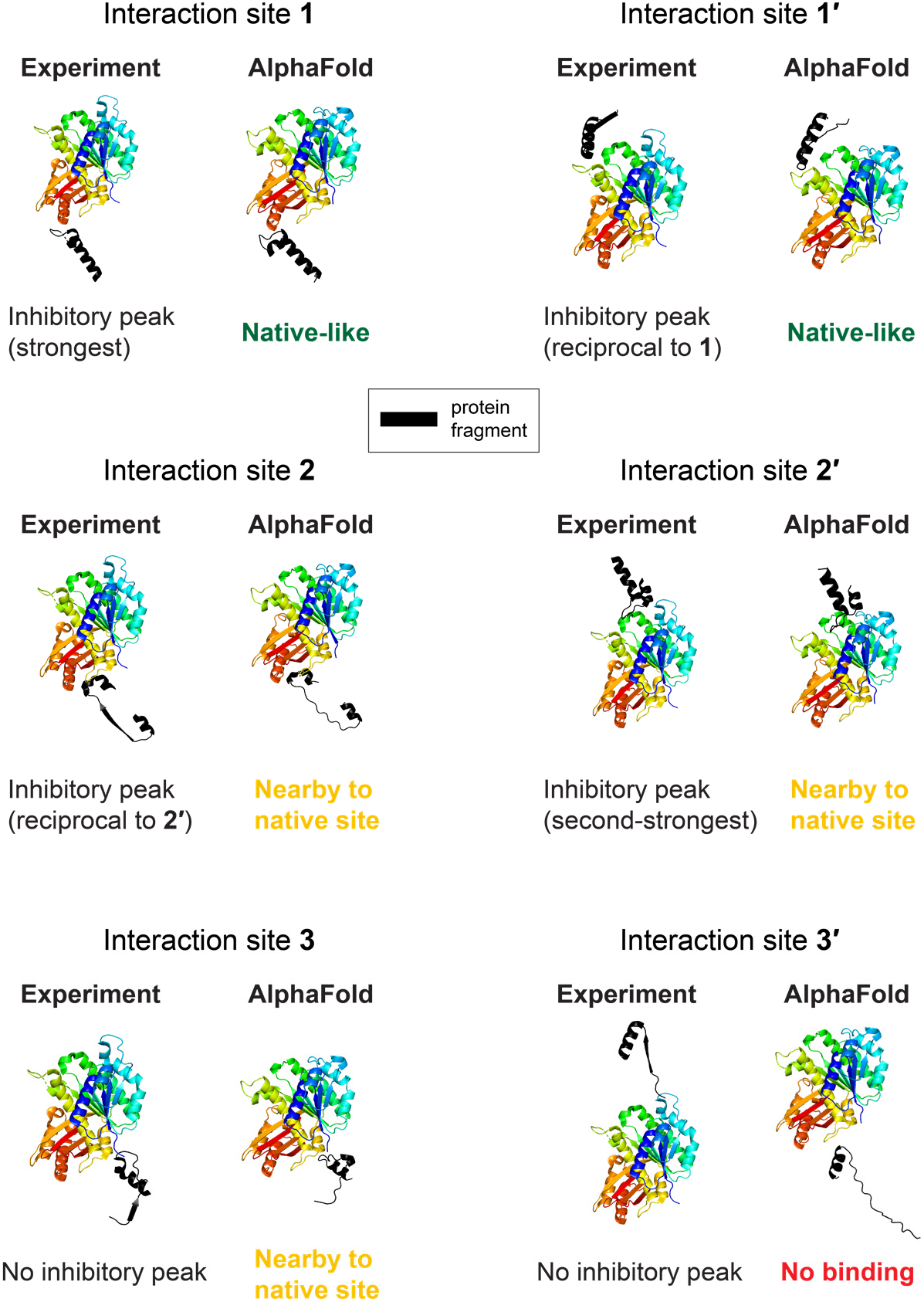
Computational modeling of FtsZ protein-peptide complexes suggests that AlphaFold can predict strong inhibitory peaks. For each of the major interaction sites of FtsZ filament formation, **1**, **1′**, **2**, **2′**, **3**, and **3′**, a side-by-side comparison is shown between (1) the 30-residue protein fragment representing the site in question, bound to the adjacent monomer, from the experimentally-determined structure of the FtsZ filament (*left*; from PDB ID 6unx); and (2) the AlphaFold-predicted structure of the corresponding protein-peptide complex (*right*). FtsZ monomers are colored from blue (N-terminus) to red (C-terminus), as in Figure 2, and protein fragments are colored black. Experimental results from the protein fragment scanning experiments are noted below the crystallographic structures, and the predicted binding mode (or lack thereof) is noted below the predicted structures.

## Discussion

In this work, we uncovered numerous inhibitory fragment peaks localized to protein interaction sites, indicating that these sites are likely susceptible to perturbation by competitive binding. As the parental proteins are essential, these peak regions constitute promising target sites for the development of antimicrobials, potentially based on the peptides themselves (15, 16). Furthermore, these proteins are extremely well-conserved in sequence and structure across many bacterial strains and species, including human pathogens such as *Shigella*, *Klebsiella*, and *Salmonella* (29), thus making inhibition-prone sites derived from *E. coli* fragment scans likely to be directly translatable to other strains.

In the case of FtsZ, our results suggest that the **1**, **1′**, **2**, and **2′** interaction sites (Fig. 2A) employed in filament formation are good targets, but that the **3** and **3′** interaction sites are not. Although no approved antibiotic targets FtsZ, multiple inhibitors thought to target the cleft between the N- and C-terminal domains, the GTP-binding pocket, and the T7 loop (site **2′**) have been reported (30), and the small protein MciZ binds sites **1′** and **2′** simultaneously (31). The fragment scans of DHFR, GyrA, and Ssb similarly yielded both known antimicrobial binding sites and potential novel ones (*SI Text*).

The protein fragments characterized here bear a striking resemblance to biologically-encoded miniproteins (32, 33). Miniproteins consist of ≤50 residues and are frequently composed of simple structural motifs, such as individual alpha-helices (6), similar to many of the inhibitory protein fragments we report. These natively-produced peptides commonly regulate proteins by binding to specific sites, analogous to the mapping of inhibitory fragments to interaction surfaces. The functional similarities are in some cases direct. Fragments of *E. coli* FtsZ mapping to the **1-1′** and **2-2′** interaction site pairs tended to be inhibitory (Fig. 2*A*), presumably by competitively inhibiting the corresponding monomer-monomer interactions. In *B. subtilis*, the MciZ miniprotein performs this function by binding in a manner comparable to the inferred binding modes of interaction site **1** and **2** fragments, occluding the **1′** and **2′** (T7 loop) interaction sites and thereby inhibiting filament assembly and cell division (31). The lack of inhibitory peaks mapping to the **3** and **3′** interaction sites suggests that these are poorer sites for competitive inhibition, consistent with the MciZ binding target.

Hydrophobic alpha-helical miniproteins of <50 residues, often just long enough to traverse the membrane, are common in bacteria (4, 6, 34), and structures of such miniproteins bound to their membrane protein targets have been determined (*e.g.*, ref. (35)). These miniproteins regulate their membrane protein targets; for example, AcrZ modulates antibiotic export by the AcrAB-TolC efflux pump (6, 36). Such miniproteins readily enter the inner membrane, including through a mechanism mediated by post-translational binding to the signal recognition particle (SRP), presumably through the affinity of SRP for hydrophobic sequences (37). It seems likely, then, that protein fragments of LptG can also take advantage of this mechanism to enter the membrane. Furthermore, LptG is present in cells at a low protein copy number (Table 1), suggesting that even limited membrane entry by its inhibitory fragments may perturb lipopolysaccharide transport.

This work highlights important considerations for future applications of high-throughput inhibitory fragment assays. Parental proteins must be selected carefully based on cellular concentration; abundant proteins would require substantially higher levels of peptide to inhibit their interactions. Protein fragment length is another crucial variable that must be controlled; for example, inhibition by fragments mapping to different sequence regions can be directly compared only if they are of the same size. In applications for which the primary goal is peptide inhibitor development, longer fragments (here ∼30-residue) are preferred as these provide more robust inhibitory activities. In studies geared towards functional mapping of interaction sites, scanning with multiple peptide fragment lengths (*e.g.*, 14 to 30 residues) might be best, as shorter fragments yielded higher-resolution maps while longer fragments more readily identified weaker peaks. Fragments that are too long (here ∼50 residues) should generally be avoided due to increased background inhibition and loss of site-specific inhibition.

Given its effectiveness in *S. cerevisiae*, *H. sapiens*, and now *E. coli* cells, inhibitory fragment scanning should be generalizable to essentially any genetically tractable species, enabling comparative analyses of fragment inhibition across orthologous proteins. Total proteome-wide fragment mapping is also within reach, though currently limited to a lower resolution than the measurements in this work. This approach also holds promise for comprehensively measuring the relative importance of interaction sites under multiple environmental settings and drug treatments, enabling the elucidation of condition-specific roles of interactions across the proteome.

## Materials and Methods

### Tiling fragment library construction

Tiling oligonucleotides were designed to cover each parental protein in each fragment size of interest with a 3-bp (one codon) step size. DNA oligonucleotides encoding these fragments were array-synthesized by Twist. Oligo sequences were centered around an exact match to the desired region of endogenous parental protein sequence, with the following features added in the flanking sequences of each fragment: (1) flanking sequences for Gibson assembly into the desired plasmid vector; (2) gene-specific pairs of 3nt indices at the 5′ and 3′ ends of the fragment, which uniquely identified the parental protein and allow selective PCR amplification from the library (Table S2); (3) a stop codon at the 3′ end of the fragment (downstream of the gene-specific 3bp index). Array-synthesized ssDNA libraries were amplified into dsDNA by qPCR following manufacturer recommendations. The resulting dsDNA libraries were cloned into the multiple cloning site of the pET-9a expression vector (Novagen) by Gibson assembly (NEBuilder HiFi DNA Assembly kit, NEB). Fragment-encoding sequences were expressed under control of a T7 promoter in a shared exogenous sequence context provided by the vector. The pET-9a vector adds an N-terminal T7 tag peptide and short linker to expressed polypeptides, so the resulting sequence of each protein fragment had the following structure: <u>MASMTGGQQMG</u>RGS**-**X1-(**fragment**)-X2**\*** where the T7 tag sequence is underlined, and X1 and X2 are serine, alanine, or leucine, encoded by the gene-specific 3bp indices.

**Table 2:** Overview of highlighted inhibitory peaks. Summary of example inhibitory peaks highlighted in the text, for each *E. coli* protein assayed by inhibitory fragment scanning.

| *E. coli* Protein | Example inhibitory peak locations in sequence (aa) | Notes |
| --- | --- | --- |
| DHFR | 17, 120 | Peaks map to a substrate-binding region (17) and a known refolding-inhibitory region (120) |
| GyrA | 386, 493 | Strongest two peaks both map to dimeric C-gate which controls DNA strand transfer directionality; one peak (386) maps to the homodimer contact required for gate closure, the other (493) to the C-gate “hinge” structure |
| IleS | Cluster of peaks at positions 600 – 800 | Example peaks map within a known refolding-inhibitory region |
| FtsZ | 131, 178, 201, 273, 368 | Example peaks map to reciprocal interaction sites for filament formation (131 and 201; 178 and 273) and an intrinsically disordered region implicated in regulatory interactions with other proteins (368) |
| RpoB | 211 | Strongest peak maps to a protein region that interacts with ssDNA |
| RplL | 20 | Peak maps to a region of RplL (50S L7/L12) involved in homodimerization and interaction with L10, required for ribosome stalk formation |
| Ssb | 67 | Peak maps to only region of defined secondary structure, involved in Ssb tetramer formation |
| GroEL | 18, 102, 176, 244/261, 446, 501 | Multiple peaks map to the GroEL-GroEL heptameric ring interface (18, 102, 446, and 501) |
| GroES | — | No inhibitory peaks observed |
| LptG | 21, 77, 292, 324 | All four peaks map to specific transmembrane domains (TM1, TM2, TM4, TM5) |

Two tiling fragment-encoding libraries were constructed in this manner: one containing 20- and 30-residue scans of the cytosolic proteins investigated (Library 1), and another containing 50-residue scans of cytosolic proteins and 14-residue scans of the membrane protein LptG (Library 2). Each library contained ∼6000 fragments. Thus, Library 2 contained a larger proportion of longer protein fragments, which are more inhibitory on average. This compositional difference between the libraries might produce systematic differences in growth rates that affect the relative inhibitory peak depths with 50-residue fragments, compared to 20- and 30-residue fragments (Fig. 4C). However, a set of 25-residue fragments derived from RplL and eGFP that were common to both libraries generally gave highly similar results (Fig. S5), yielding an average enrichment score difference of 0.08 (frequency change fold-difference of 1.06) with a standard deviation of 0.57 (frequency change fold-difference range of 0.71 – 1.58).

Following Gibson assembly, tiling fragment libraries were transformed into ElectroMax *E. coli* cells (Thermo Fisher). Plasmids were isolated from the transformed ElectroMax cells (Qiaprep Spin Miniprep Kit, Qiagen), and assembled library composition was confirmed by high-throughput sequencing. These purified plasmid libraries served as the starting point for subsequent selection experiments.

### Massively-parallel measurements of dominant-negative inhibition by protein fragments

The inhibitory activity of protein fragments was measured in a high-throughput manner via growth selection experiments, similarly to previous studies (14, 15). Tiling fragment-encoding plasmid libraries were transformed into highly electrocompetent *E. coli* BL21(DE3) cells (Sigma-Aldrich), yielding transformants at >200-fold coverage of the library size. After 1 hr in recovery medium, each 1 mL of transformed cells was transferred to 50 mL of LB medium + 50 µg/mL kanamycin + 10 µM IPTG to begin the growth selection. These cultures were grown at 37 °C with shaking at 220 rpm. Multiple replicate experiments, entailing independent transformations and growth selections, were performed for each library (*N* = 4 for Library 2 experiments with 20- and 30-residue fragments, and *N* = 2 for Library 1 experiments with 14- and 50-residue fragments). Cells were grown to a final OD_600_ of 0.6 – 0.9 and harvested by centrifugation; this constituted the endpoint of the selection. Plasmids were isolated from each sample by miniprep (Qiagen). The selection endpoint samples and the plasmid library input were prepared for sequencing as follows.

PCR amplification (≤12 cycles) was used to extract fragment-coding sequences from the plasmid library and add sequencing adapters and sample-identifying indices. High-throughput sequencing was performed using Illumina’s NextSeq 550 platform. Paired-end sequencing was employed to uniquely identify fragments by their 5′ and 3′ ends. Sequencing was performed with ∼6 million reads per sample (1000-fold coverage).

### Fragment assay data analysis

The identity (parental protein, sequence location, and orientation) of each fragment was determined by aligning each read pair to the set of gene sequences included in the library using Bowtie2. Fragment counts in each sample were determined, and fragments with insufficient read depth in the input sample were filtered out of the dataset (<50 reads for Library 1, and <100 reads for Library 2). Fragment sequences that completely dropped out in the growth selection were assigned pseudocounts of 0.5 in the corresponding output samples. Fragment frequencies in each sample were then determined from the counts of each fragment in each sample, divided by the total counts of all fragment sequences in the sample. The enrichment (*E*) of each fragment was determined from its growth selection input and output frequencies as *E* = log2(*f*post-selection/*f*pre-selection), where *f* denotes frequency. *E* < 0 therefore indicates depletion. Mean and s.e.m. values of *E* were calculated for each fragment based on multiple replicate experiments. Fragment maps were generated by plotting *E* for each fragment tiling a protein as a function of the fragment’s position along the parental protein sequence. Average and maximum inhibitory effects of fragments derived from a protein were determined by calculating the mean and minimum of *E,* respectively, for the appropriate subsets of fragments. Parental protein structures overlaid with representative fragments or widths of inhibitory peaks were generated from protein structures found in the RSCB PDB using Pymol.

Cellular concentrations of parental proteins were retrieved from the data of Li and colleagues (24), as maintained by the EcoCyc database (25); the protein concentrations determined under rich media conditions (Neidhardt EZ rich defined medium) were used. Physicochemical properties of protein fragments were determined using the Peptides package in R (38). In particular, Peptides was employed to calculate the peptide properties hydrophobicity (Kyte-Doolittle scale (39)); charge (Lehninger pKa scale (40), at pH 7); and instability (Guruprasad instability index (27)). For the ANOVA analysis, each fragment was classified as “hydrophobic” if the Kyte-Doolittle hydrophobicity was > 0, and “hydrophilic” otherwise; as “neutral” if the Lehninger pKA-based charge was in the range (–1,1), “negative” if charge < –1, and “positive” if charge > 1; and as “unstable” if the Guruprasad instability index was ≥40 (following the standard definition (27)), and “stable” otherwise. Alpha-helical, beta-strand, and turn content of protein fragments in the full-length protein context were determined as follows. Secondary structure annotations were retrieved from each parental protein’s Uniprot entry (41), and each protein fragment was annotated as containing each type of secondary structure content if the sequence region covered by the fragment overlapped with the corresponding secondary structure feature; otherwise the fragment was annotated as lacking that type of secondary structure.

N-way nested analysis of variance (ANOVA) was performed using the built-in “anovan” function in Matlab. The following parameters were included in the analysis: fragment length; parental protein; hydrophobicity, charge, and predicted instability, classified as described above; fragment alpha-helical, beta-strand, and turn content (a binary True/False classification for each); and fragment relative position in the parental protein sequence (classified as “N-terminal” for fragments sourced from the first third of the sequence; “central” for fragments sourced from the central third; and “C-terminal” for fragments sourced from the C-terminal third); finally, fragment identity (a unique tag assigned to each fragment based on parental protein, sequence start and stop sites, and orientation) was nested under all other parameters. The measurable variable was fragment enrichment, *E*. ANOVA was performed on the combined data from both tiling libraries, excluding only fragments of IleS, due to a lack of structural information which prevented assignment of fragment secondary structure content.

### DHFR random fragment library experiments

The random fragment library of DHFR was generated as previously described (14). Briefly, the *folA* gene was PCR-amplified out of the *E. coli* genome and fragmented using the Nextera transposase (Illumina). The resulting gene fragments were amplified by PCR to add sequence adapters for Gibson assembly (as well as stop codons in all three frames), and cloned into the pET-9a vector backbone at the multiple cloning site. Growth selection experiments were performed as described for the tiling fragment libraries, except 100 µM IPTG was used. High-throughput sequencing of input and output library samples and subsequent analyses to determine fragment localization, frequencies, and enrichment were performed as for the tiling libraries. Average per-site enrichment was calculated by determining the average *E* of all fragments whose sequence covers a given position in the parental protein sequence.

### AlphaFold computational predictions of peptide-protein complexes

Peptide-protein complexes were modeled using Alphafold2 (28), employing the AlphaFold2_complexes notebook hosted on the Google Colab servers (42, 43). Input query sequences for each model were (1) the full-length sequence of *E. coli* BL21(DE3) FtsZ and (2) the sequence of a 30-residue fragment of FtsZ representing an inhibitory peak or other sequence region of interest. Default settings were used for other parameters: num_models = 5, msa_mode = MMseqs2, pair_msa = False. The structures reported in the paper correspond to output model 1 from each prediction.

### Author Contributions

Research conceptualization and design were performed by A.S. and S.F. DHFR random fragment library construction and preliminary selection experiments with this library were performed by A.S. and A.F. All other experiments were performed by A.S. Data analysis was performed by A.S. The manuscript was written by A.S. and S.F.

## Acknowledgements

We thank R.M. Garner, G.W. Li, G. Storz, D.K. Schweppe, A.E. Keating, C.D. Bahl, M.W. Dorrity, B.M. Brandsen, and members of the Fields and Li labs for helpful discussions. We are grateful to M.W. Dorrity, W.J. Greenleaf, C. Queitsch, and G.W. Li for comments on the manuscript. A.S. was supported by NRSA fellowship F32 GM134557 from the National Institutes of Health. This work was supported by grants RM1 HG010461 and P41 GM103533 from the National Institutes of Health.

## Supplementary Information

### Supplementary Figures

**Figure S1:**
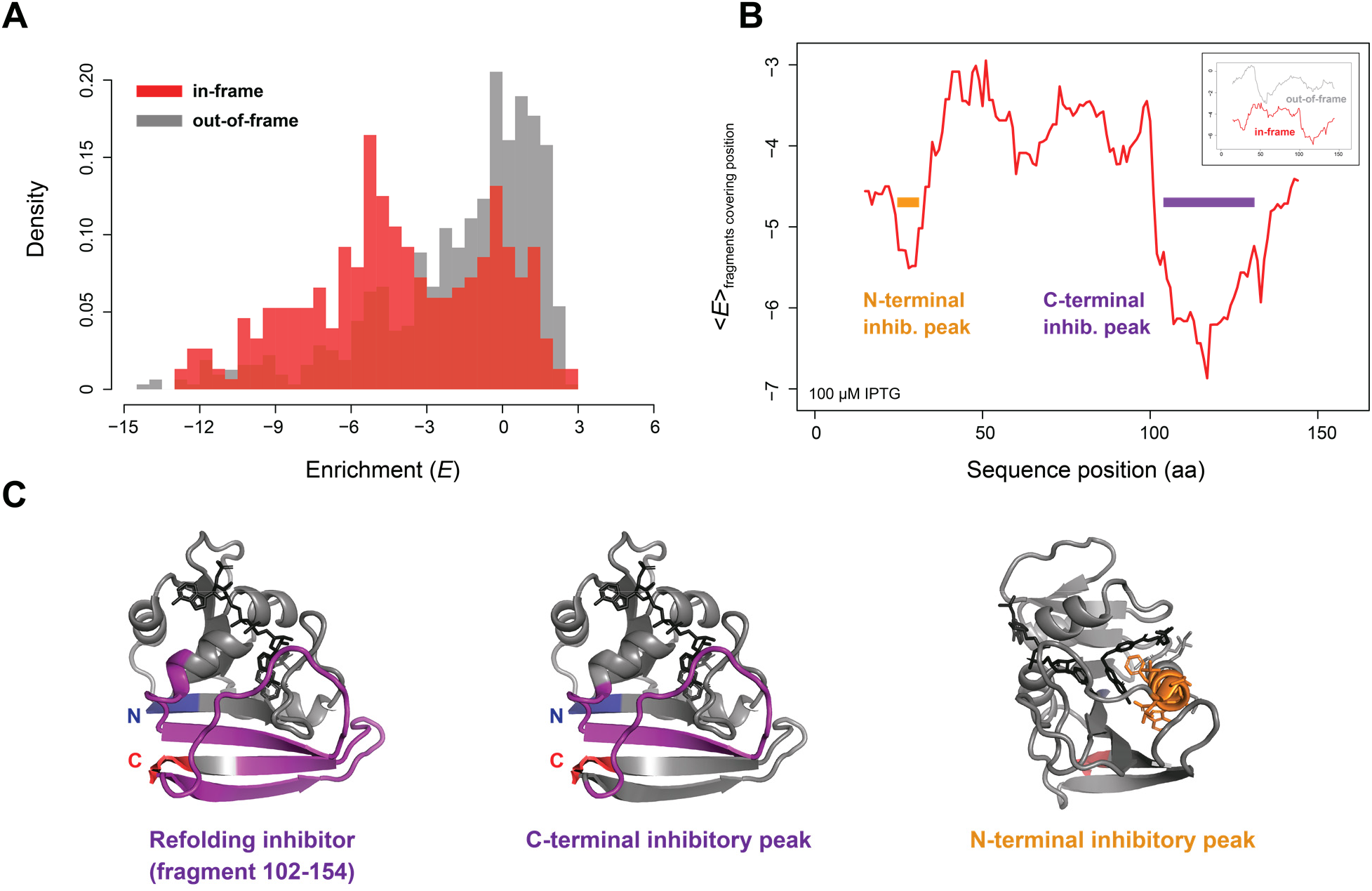
A random protein fragment assay of DHFR shows that inhibition is reading frame-dependent. (*A*) Histogram of enrichment (*E*) by in-frame (red) and out-of-frame (grey) fragments of ≤30 residues derived from the random fragmentation of *E. coli* DHFR. (*B*) Average enrichment per site due to in-frame fragments covering each residue, across the DHFR sequence (see *Materials and Methods*). Inhibitory peaks are indicated. Inset: overlay of per-site enrichment of in-frame and out-of-frame fragments. (*C*) DHFR crystallographic structure (PDB ID 7dfr) with N- and C-termini indicated, overlaid by the known refolding inhibitory fragment (*left*), alongside the C-terminal inhibitory peak (*middle*) and the N-terminal inhibitory peak (*right*) from the random fragment assay.

**Figure S2:**
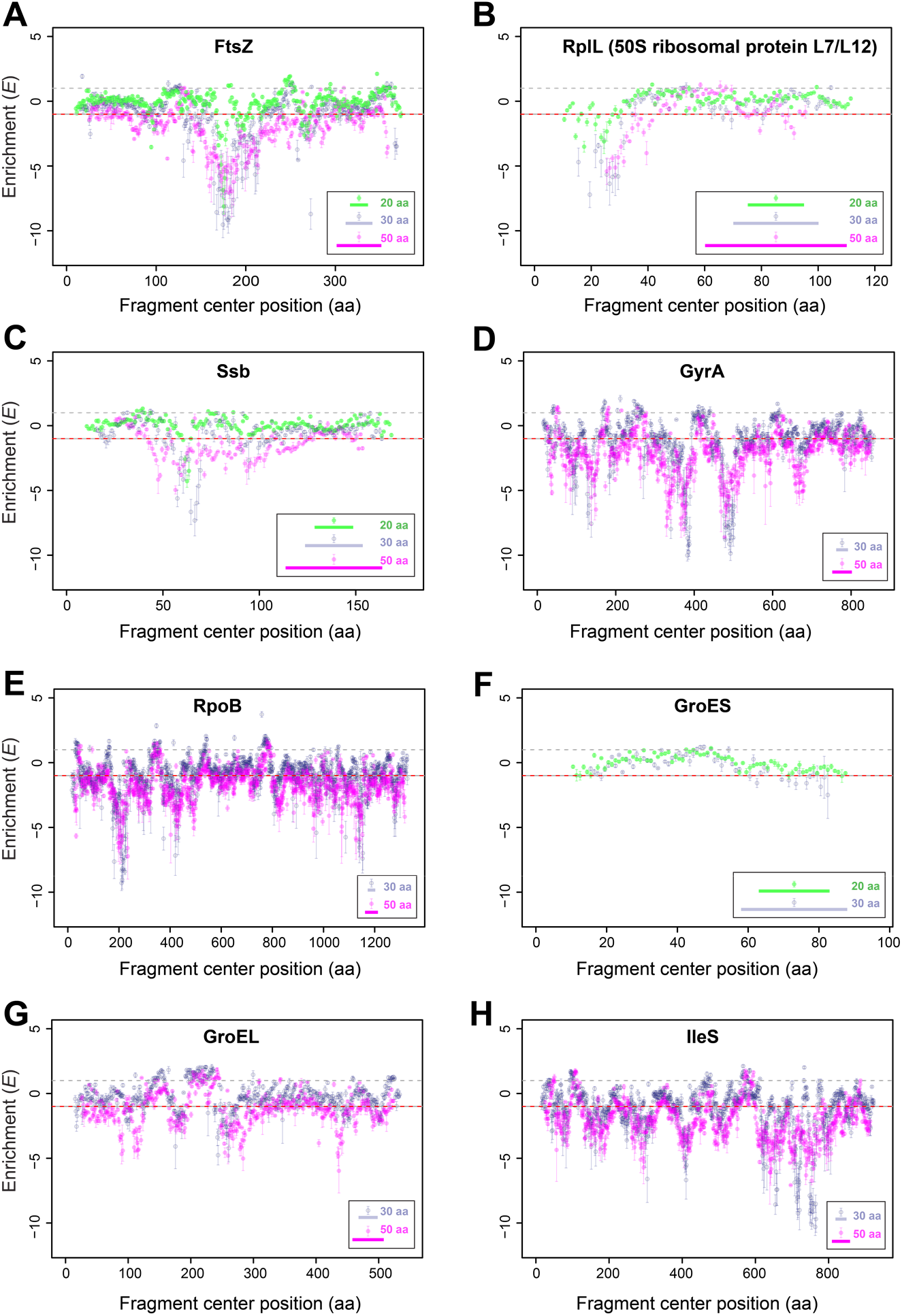
Details of fragment length effects across diverse proteins. (A) – (*H*) Inhibitory fragment scans of *E. coli* proteins with tiling fragments, at a tile step size of 1 amino acid, comparing different fragment lengths. Data are plotted as in Figure 2, with individual fragment enrichments (*E*) and their s.e.m. across multiple experiments shown, except that 30-residue fragment data are shown in blue-grey; 20-residue fragment data in green; and 50-residue fragment data in magenta.

**Figure S3:**
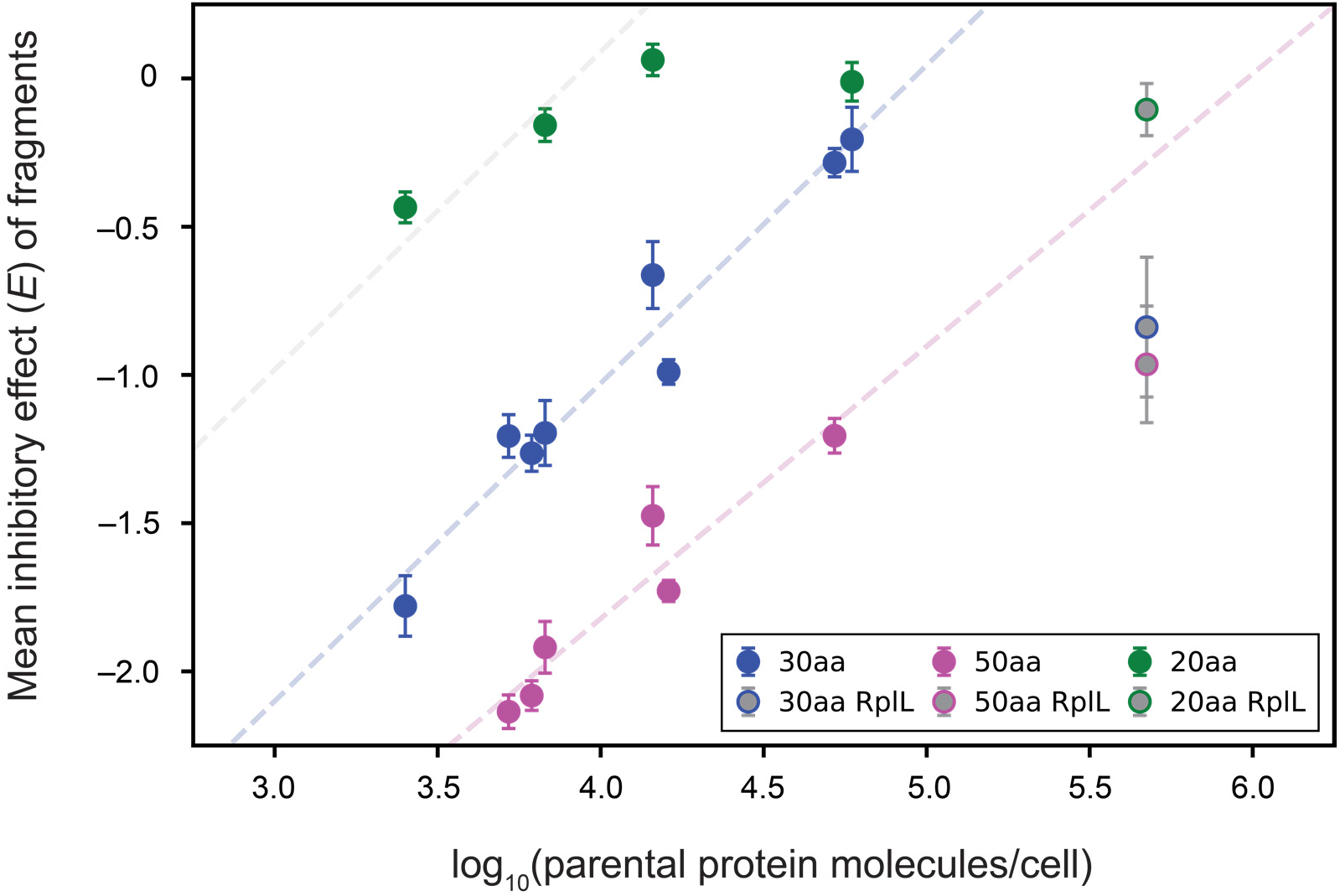
Average susceptibility to inhibition by 50-residue fragments remains negatively correlated with parental protein concentration. Mean inhibitory effects (selection enrichment *E*) of 20-, 30-, and 50-residue fragments tiling each cytosolic protein investigated, as a function of the log of the parental protein concentration. Error bars: s.e.m. Not all proteins were tiled with all fragment sizes. 50S ribosomal subunit protein L7/L12 (RplL) is shown with a grey fill color; linear fits to the non-ribosomal protein data are shown for the 30-residue and 50-residue data as dashed lines in corresponding colors. Note that the slopes are very similar. The faded grey dashed line near the 20-residue fragment data is a shifted copy of the trendline for the 30-residue data, suggesting that a similar slope may also be reasonable for 20-residue fragments; however, among the proteins for which 20-residue fragment libraries were generated, a leveling off is evident as the mean *E* enters the neutral regime (E ≈ 0), as expected, and there is insufficient sampling of mean *E* values below zero to fit these data directly.

**Figure S4:**
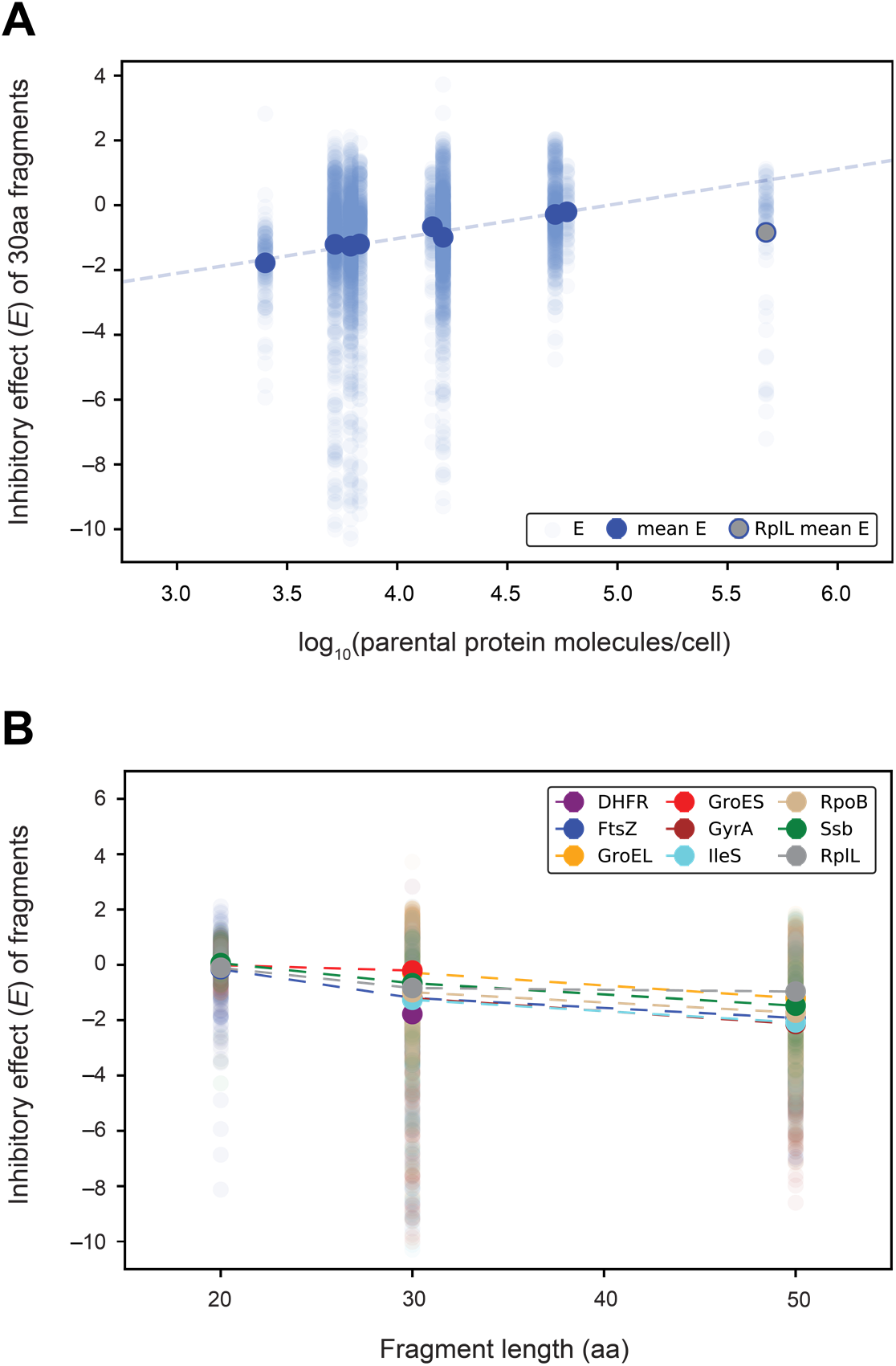
Parental protein concentration and fragment length effects explain only a fraction of individual fragment-to-fragment variability. (*A*) Mean inhibitory effects of 30-residue fragments derived from each protein investigated (large blue markers), and corresponding linear fit to the non-ribosomal proteins, as plotted in Figure 3, overlaid with the inhibitory effects (*E*) of individual fragments of each protein in pale blue. (*B*) Mean inhibitory effects of 20, 30, and 50-residue fragments of each protein, as in Figure 4*B* (large markers connected by dashed lines), overlaid with the inhibitory effects (*E*) of individual fragments (smaller, pale markers). Error bars indicating s.e.m. are included for the protein averages but are smaller than the markers.

**Figure S5:**
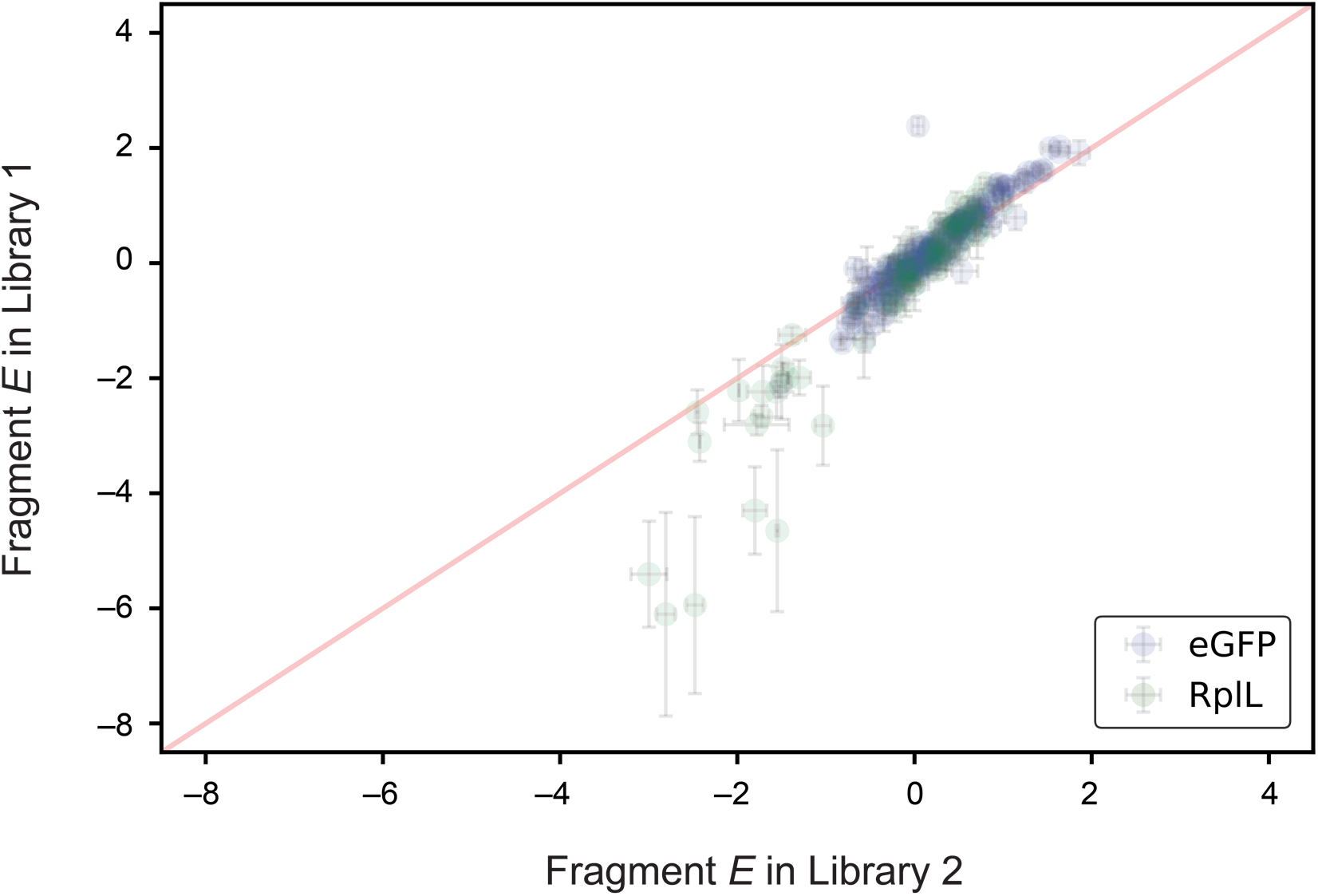
Comparison of shared fragment enrichments in the two tiling fragment libraries. Enrichment (*E*) of a set of 25-residue fragments of eGFP and RplL shared between the two tiling fragment libraries (*Materials and Methods*). The red line indicates *y* = *x*. Error bars: s.e.m. Note that points with lower measurement error tend to be highly reproducible between libraries, and the points farther from the diagonal generally exhibit high measurement error.

### Supplementary Tables

**Table S1:** Results from analysis of variance performed on fragment-based inhibition data.

| Source of Variation | Sum of Squared Errors | Fraction of Total Variation (%) | d.f. | P-value |
| --- | --- | --- | --- | --- |
| **Fragment ID** | 17371.55 | 71.6946 | 9921 | 2.97E-93 |
| **Fragment length, aa** | 2419.80 | 9.9868 | 40 | 1.46E-227 |
| **Parental protein** | 1021.78 | 4.2170 | 9 | 2.30E-191 |
| **Fragment contains beta-strand (from structure)** | 173.34 | 0.7154 | 1 | 1.88E-97 |
| **Fragment contains alpha-helix (from structure)** | 118.07 | 0.4873 | 1 | 4.18E-79 |
| **Fragment contains turn (from structure)** | 71.92 | 0.2968 | 1 | 1.87E-58 |
| **Hydrophobicity (Kyte-Doolittle)** | 169.05 | 0.6977 | 1 | 3.39E-96 |
| **Relative position in sequence (N-->C)** | 53.77 | 0.2219 | 2 | 6.29E-47 |
| **Instability (Guruprasad)** | 2.54 | 0.0105 | 1 | 1.57E-04 |
| **Net charge (Lehninger)** | 0.56 | 0.0023 | 2 | 2.01E-01 |
| *Error* | *52.08* | *0.2150* | *300* | — |

**Table S2:** Table of single-codon indices appended to protein fragments in the tiling library.

| **Protein** | **N-terminal index**<br>**codon (amino acid), X<sub>1</sub>** | **C-terminal index**<br>**codon (amino acid), X<sub>2</sub>** |
| --- | --- | --- |
| **DHFR** | gca (A) | ctg (L) |
| **GyrA** | gca (A) | agt (S) |
| **IleS** | gca (A) | gca (A) |
| **FtsZ** | ctg (L) | ctg (L) |
| **RpoB** | ctg (L) | gca (A) |
| **RplL** | ctg (L) | agt (S) |
| **Ssb** | agt (S) | ctg (L) |
| **GroEL** | agt (S) | gca (A) |
| **GroES** | agt (S) | agt (S) |
| **LptG** | tcc (S) | ctg (L) |

### SI Text

#### Fragment mapping reveals promising target sites for inhibitor development: additional discussion

DHFR is the target of the antibiotic trimethoprim, which binds at the enzymatic active site and inhibits catalysis (44). Our results suggest that inhibition of folding by compounds that bind to the N-terminal beta strand may be another approach to target this important enzyme. Another route would be welcome given the serious issue of trimethoprim resistance (45); due to the distinct mechanisms, a putative folding inhibitor would likely not be subject to the same resistance mutations.

A number of compounds inhibiting bacterial DNA gyrase, including the fluoroquinolones currently employed clinically, function by trapping the enzyme in a DNA cleavage complex (46). Other compounds, such as the aminocoumarins, interfere with ATPase activity (46). Our finding of an inhibitory fragment peak localized to the ends of the C-gate arms of GyrA, which form a homomeric head-to-head complex to close the gate, suggests that compounds targeting these C-gate closure sites would provide another means to inhibit gyrase. Such inhibitors would rely on a distinct binding mode from the aforementioned compounds, which bind at sites at or near the DNA gate or the ATPase site (46, 47). In line with our findings, a monoclonal antibody against *Mycobacterium smegmatis* and *Mycobacterium tuberculosis* GyrA appears to bind at the same location as our fragments and prevent C-gate dimerization. Furthermore, the antibody epitope corresponds to *E. coli* GyrA residues 340-402 (48), fully overlapping with the inhibitory fragment peak centered around residue 386.

Single-stranded DNA binding protein (Ssb) is a desirable antibiotic target because inhibition should interfere with DNA replication and repair. Screening has therefore been performed for Ssb inhibitors (49–51), leading to the identification of numerous compounds; some of these were shown to interfere with critical interactions of eight conserved C-terminal residues with genome maintenance proteins (50, 51). Unlike the case for FtsZ, we did not observe an inhibitory fragment peak mapping near this interaction-mediating C-terminal tail. Instead, we observed a strong peak mapping to the single alpha-helical element of this protein, suggesting a novel approach for inhibitor development: mimicking the interactions formed by the alpha helix in Ssb dimers to disrupt dimer and tetramer formation.

A further note regarding the transferability of inhibitory sites identified in *E. coli* to other bacteria: The range of bacteria potentially targetable based on these fragment scan results is extensive, given that protein-peptide interactions are based heavily on structural properties. Therefore, more distantly related orthologs retaining the relevant structural features are likely susceptible to the same types of inhibitors, especially those based directly on peptide fragments.

